# Discovery, Expression, Cellular Localization, and Molecular Properties of a Novel, Alternative Spliced HP1γ Isoform, Lacking the Chromoshadow Domain

**DOI:** 10.1101/638395

**Authors:** Angela Mathison, Thiago Milech De Assuncao, Nikita R. Dsouza, Monique Williams, Michael T. Zimmermann, Gwen Lomberk, Raul Urrutia

## Abstract

By reading the H3K9Me3 mark through their N-terminal chromodomain (CD), HP1 proteins play a significant role in cancer-associated processes, including cell proliferation, differentiation, chromosomal stability, and DNA repair. Here, we used a combination of bioinformatics-based methodologies, as well as experimentally-derived datasets, that reveal the existence of a novel short HP1γ (CBX3) isoform, named here sHP1γ, generated by alternative splicing of the *CBX3* locus. The sHP1γ mRNA encodes a protein composed of 101 residues and lacks the C-terminal chromoshadow domain (CSD) that is required for dimerization and heterodimerization in the previously described 183 a. a HP1γ protein. Fold recognition, order-to-disorder calculations, threading, homology-based molecular modeling, docking, and molecular dynamic simulations show that the sHP1γ is comprised of a CD flanked by intrinsically disordered regions (IDRs) with an IDR-CD-IDR domain organization and likely retains the ability to bind to the H3K9Me3. Both qPCR analyses and mRNA-seq data derived from large-scale studies confirmed that sHP1γ mRNA is expressed in the majority of human tissues at approximately constant ratios with the chromoshadow domain containing isoform. However, sHP1γ mRNA levels appear to be dysregulated in different cancer types. Thus, our data supports the notion that, due to the existence of functionally different isoforms, the regulation of HP1γ-mediated functions is more complex than previously anticipated.

## Introduction

HP1 is a cancer-associated chromatin protein, which was first identified as a major component of heterochromatin[1, 2]. HP1 is highly conserved from *S. pombe* to mammals, in which three closely related paralogs exist: HP1α, HP1β, and HP1γ[3]. Three HP1 family members in mammals are similar in amino acid sequences and structural organization, but functionally distinct. The structure of HP1 proteins includes an N-terminal chromodomain (CD), followed by a linker region, and then a C-terminal chromoshadow domain (CSD)[4]. HP1 is an elongated molecule in which intrinsically disordered regions (IDRs) allow this protein to have dynamic flexibility, intermolecular recognition properties, and the ability to integrate signals from various intracellular pathways. Extensive structural and biochemical studies have demonstrated that this modular structure is of paramount importance for HP1 proteins to perform their molecular and cellular functions[4–8]. Indeed, HP1 proteins use their CD to bind either the di-methylated or tri-methylated forms of H3K9 within genomic regions that are marked in this manner to be transcriptionally silent[5, 6]. The linker region, which lies in between the CD and CSD domains, contains a nuclear localization signal (NLS) and a DNA binding motif[4, 7, 8]. Finally, the CSD is used by HP1 proteins to homo- or hetero-dimerize among themselves, which is the most common manner to regulate their functional specificity. Notably, CSD dimerization leads to the formation of a docking motif, which is used by HP1 proteins to interact with large variety of chromatin regulators, typically through a PXVXL consensus motif[4]. These additional interactions allow the execution of complex-specific cancer-associated functions, including DNA recombination and repair, cell proliferation, differentiation, and migration, as well as chromosomal stability[4, 9]. Interestingly, studies on HP1 have stimulated the field of epigenomics in a manner that has resulted in the discovery of a larger family of proteins, which, by having similar domains to HP1, were found to display related functions in chromatin remodeling and epigenomics. By analogy to HP1, these CD-containing proteins are collectively known as Chromobox proteins, or CBXs, and are primarily involved in gene silencing[10]. Thus, the significant number of biochemical and biophysical studies that have been focused on defining the structure-function relationship of the different domains within HP1 have advanced our understanding of one of the most important families of proteins involved in epigenomic regulation.

Our laboratory studies the role of HP1-related pathways in pancreatic ductal adenocarcinoma (PDAC), since several members of the pathway are deregulated in this type of aggressive tumor and others[9, 11]. Surprisingly, during the course of RNA-Seq experiments[12], we discovered that the *CBX3* locus, which encodes for HP1γ, undergoes alternative splicing, leading to the generation of the conventional CD-linker-CSD containing HP1γ protein, as well as a novel shorter isoform (sHP1γ). Using qPCR and an anti-sHP1γ antibody, we confirm that, indeed, this spliced isoform is expressed in a variety of tissues and retains the typical nuclear localization expected from its functions. However, while this shorter isoform preserves a CD, it lacks the CSD contained within the conventional longer isoform. Thus, we infer that alternative splicing of the *CBX3* locus gives rise to different proteins, which have distinct structural and molecular properties to likely function in both overlapping and divergent manners.

## Results

### Identification of sHP1γ reveals an expanded repertoire of HP1γ isoforms in human tissues

Our laboratory has made significant contributions to the study of HP1 proteins in transcriptional regulation, epigenetics and cancer. In fact, the current study was derived from initial experiments looking at the expression pattern of HP1 proteins in pancreatic cancer cells. Careful examination of the reads obtained by RNA-Seq[12] suggested that the *CBX3/HP1γ* locus undergoes alternative splicing to give rise to a new protein-coding isoform. The *CBX3* locus is located within human chromosome 7 and has 6 exons. *CBX3* is currently known to produce a protein-coding transcript 2128bp in length corresponding to a protein of 183 residues. The protein structure follows the typical IDR-CD-IDR-CDS-IDR structure known for all HP1s[4]. Notably, however, in this study, we report the existence of an additional 1717bp transcript which encodes a smaller protein of 101 amino acids that lacks the C-terminal CSD (Figs 1A and 1B), hereafter named short HP1γ (sHP1γ). sHP1γ is generated by a splicing event that skips part of the fourth exon, shifts the reading frame and thus initiates a new stop codon in exon 5 (Fig 1C). Thus, the CSD found at the C-terminus of the conventional HP1γ is absent in the short isoform described here, which makes this protein unique among all members of this family (Figs 1A and 1B). Additional analysis of the primary structure of this protein, using the ProtParam algorithm[13] yields an expected Mw of 11.85796 KDa and a theoretical pI of 9.63, the latter being similar to most nuclear proteins, in particular histones and those which associate to them. Additional key characteristics, calculated from the sHP1γ primary structure, are its extinction coefficient, which informs our ability to measure the concentrations of this protein, of 8480 M^−1^cm^−1^, at 280 nm measured in water, which corresponds to an absorbance of 0.715 at a concentration of 1 g/L. Regarding the amino acid composition, we find a high content of certain types of residues, which are expected to contribute to its folding pattern and functional properties. These residues include a relative high content of aliphatic and aromatic residues (12 L and 10 V, 4 A, 5 M, 4 F, 1 W, and 3 Y), which likely participate in creating the hydrophobic moment that is necessary for proteins to fold[14], as well as forming the aromatic cage, which is used by chromodomains to bind H3K9Me2 and H3K9Me3[15]. The presence of key basic residues, including 19 K and 4 R, is found in nuclear proteins, for which they form part of nuclear localization signals, as well as sites for acetylation, ubiquitination, and methylation; likely events involved in signaling within the cell nucleus. Phosphorylatable residues include 6 T, 2 S, and 2 Y. This data is in agreement with previous observations by our laboratory and others, which show that these residues are amenable to extensive modifications in HP1 proteins, working as a subcode in regulating the function of reader proteins that operate to fine-tune the histone code[16, 17]. The absence of the a CSD indicates that the protein is not able to form the expected CSD-to-CSD interactions that support homo- and hetero-dimerization of conventional HP1 proteins. Lastly, the protein has 13 E, which are known to be characteristic of CD-containing proteins likely for reinforcing their binding to histone tails that are in essence highly basic in nature[18]. Therefore, the new alternatively spliced mRNA, first identified from pancreatic cancer RNA-Seq data[12], as well as its predicted protein product would expand the number of HP1 isoforms in human tissues.

**Fig 1.**
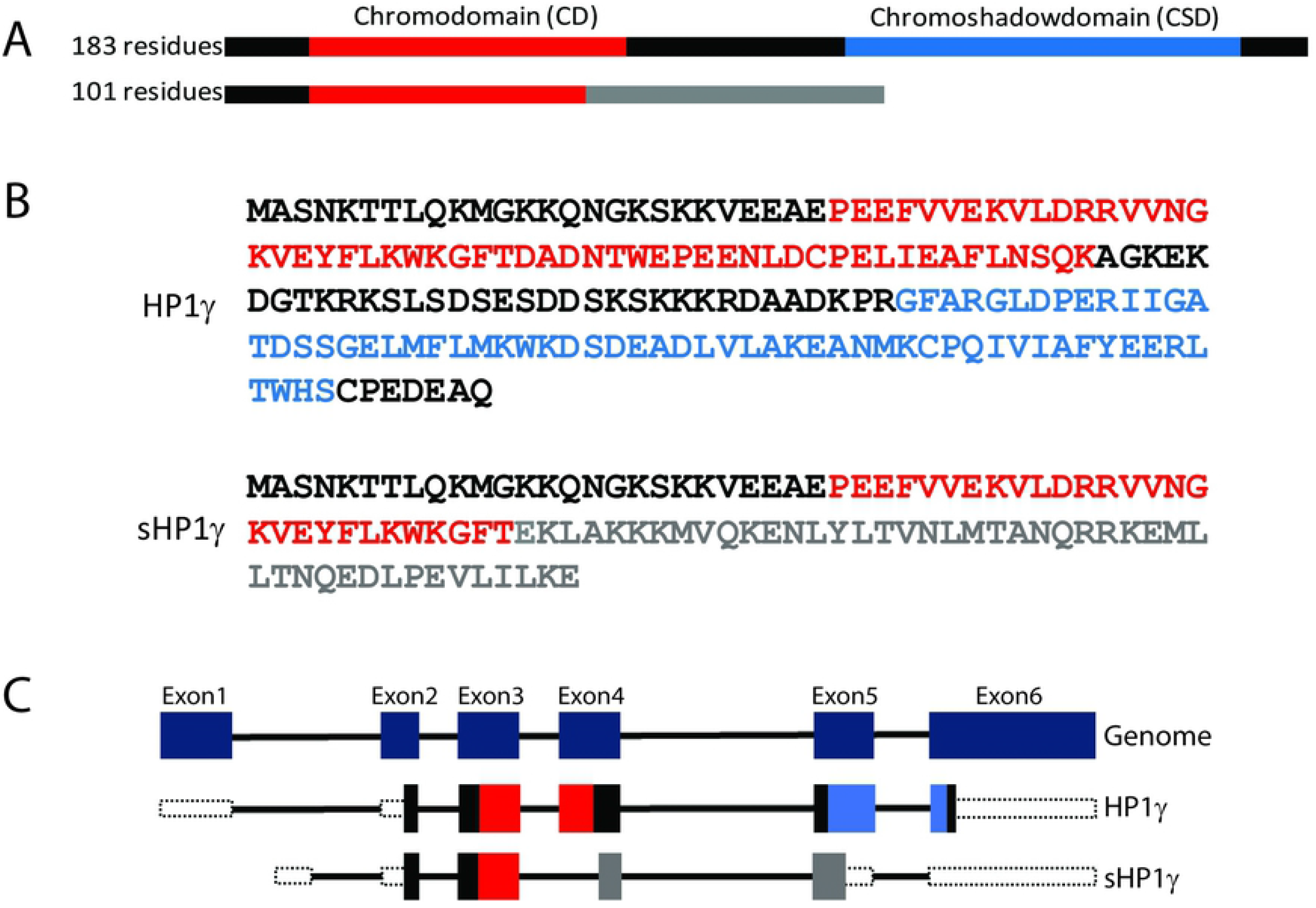
Identification of sHP1γ and comparison with canonical HP1γ. (A) Physical map of a new HP1γ isoform generated by alternative splicing. The conventional HP1γ is 183 residues and contains a chromodomain (CD) and a chromoshadow domain (CSD). A shorter isoform lacks the CSD and is 101 residues. (B) Protein sequences for HP1γ generated by alternative splicing. Sequences with highlighted CD (red) and CSD (blue), the sHP1γ sequence generated by alternative splicing is represented in gray. (C) The HP1γ genomic structure (top) with dark blue boxes representing constitutive exons. The lower panel illustrates mRNA splicing for the canonical and sHP1γ isoforms. CD is represented in red boxes, CSD is represented in blue boxes and sHP1γ sequence generated by alternative splicing is represented in gray boxes.

Subsequently, we investigated the tissue distribution of sHP1γ mRNA. For this purpose, we first used an isoform-specific qPCR method, designed to specifically detect the two different splice junctions between exon 3 and 4 that are unique to the long and short HP1γ isoforms. Using this method, we compared the expression levels of the novel sHP1γ encoding mRNA with that of the conventional HP1γ isoform (Fig 2A), in a normal human tissue mRNA panel. The results of this experiment demonstrated that both types of mRNAs are expressed in the same pattern in most human tissues. We found high levels of expression in the gastrointestinal and reproductive tracts, namely pancreas, intestine, placenta, spleen and testis, and low signals present in urinary organs, such as bladder and kidney, as well as in striated and cardiac muscle. The same isoform comparison was made among 8 commonly-used pancreatic cancer cell lines (Fig 2B), in which we detect comparable levels to those found in the normal organ. Combined, these results demonstrate that the *CBX3* locus is spliced to give rise to a small isoform, sHP1γ, in various tissues.

**Fig 2.**
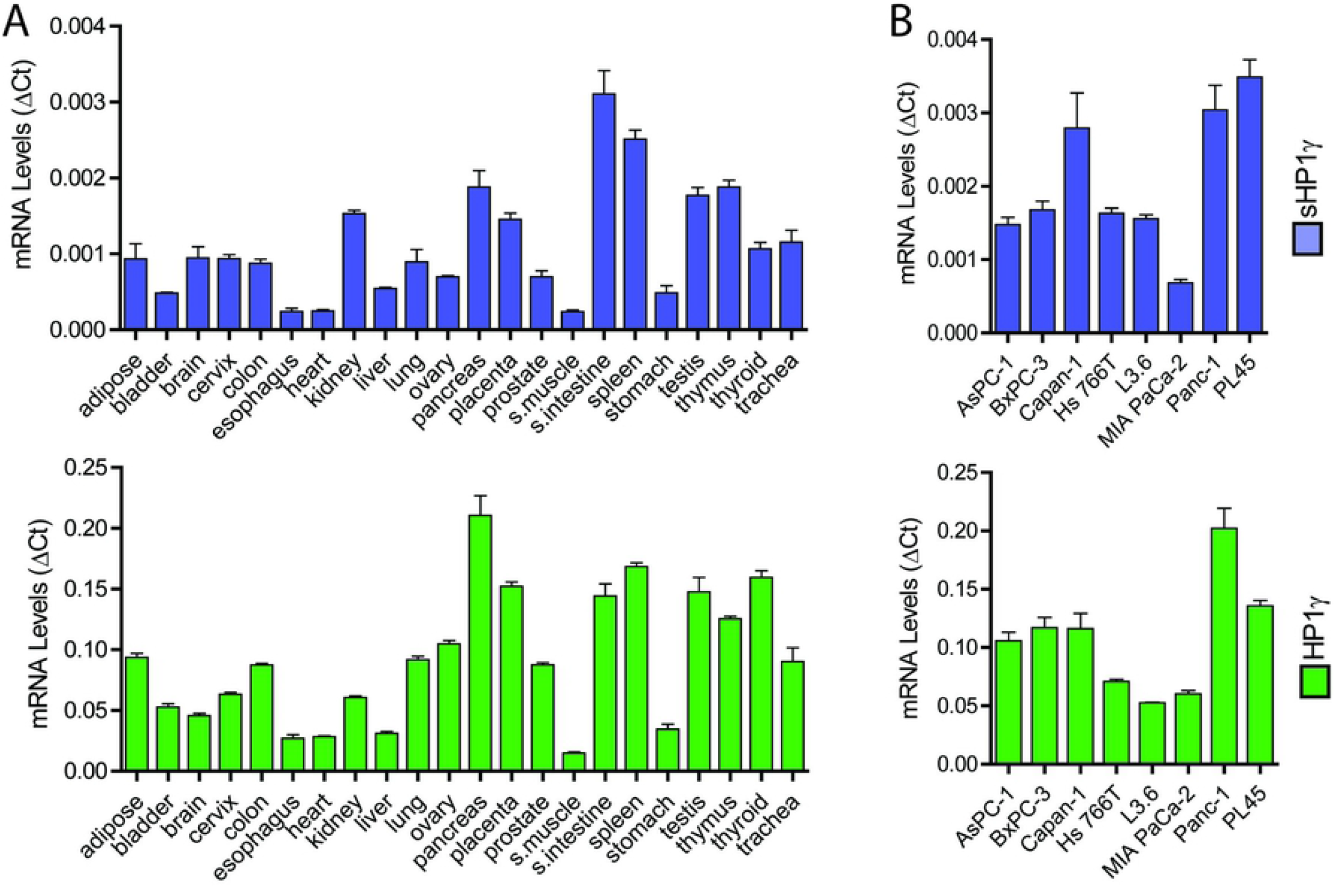
Detection of sHP1γ mRNA in Human Samples. (A) Isoform-specific qPCR assays were used to detect and compare the expression of the novel sHP1γ encoding mRNA (blue graph) with that of the long, conventional HP1γ isoforms (green graph) in 22 different human tissues. (B) The same comparison between sHP1γ (blue graph) and the conventional HP1γ (green graph) transcripts was performed in 8 pancreatic cancer cell lines.

sHP1γ transcript expression in healthy adult human tissues was obtained by processing data derived from the Genotype-Tissue Expression project (GTEx)[19]. We analyzed 53 tissues subdivided in 6 major groups (Reproductive, Other, Neurologic, Muscular, Blood, and GI). We found that sHP1γ is present in all tissues (Fig 3A) at a lower level yet in a fairly consistent ratio to the conventional HP1γ isoform (S1 Fig). Long tails on these distributions indicate that there is a small number of normal human samples that have significant up-regulation of sHP1γ, with the heatmap further emphasizing these differences (Fig 3B). We also investigated sHP1γ expression in malignant tumors using data from the Cancer Genome Atlas (TCGA) (Figs 3C and 3D) with a particular focus on defining the ratio of short to long isoform for each cancer type (S2 Fig). We noted interesting variability in the ratio of short to long HP1γ among tumors, with the highest levels of sHP1γ being found in esophageal cancer (ESCA), ovarian cancer (OV) and stomach adenocarcinoma (STAD). Compared to non-tumor tissues, tumor groups are more uniform with a much higher fraction of tumor samples from GI tissues and the blood expressing a higher level of sHP1γ. Thus, these comparisons show that the two isoforms are expressed at a consistent ratio in normal human tissues, but this ratio becomes more variable in different cancer types.

**Fig 3.**
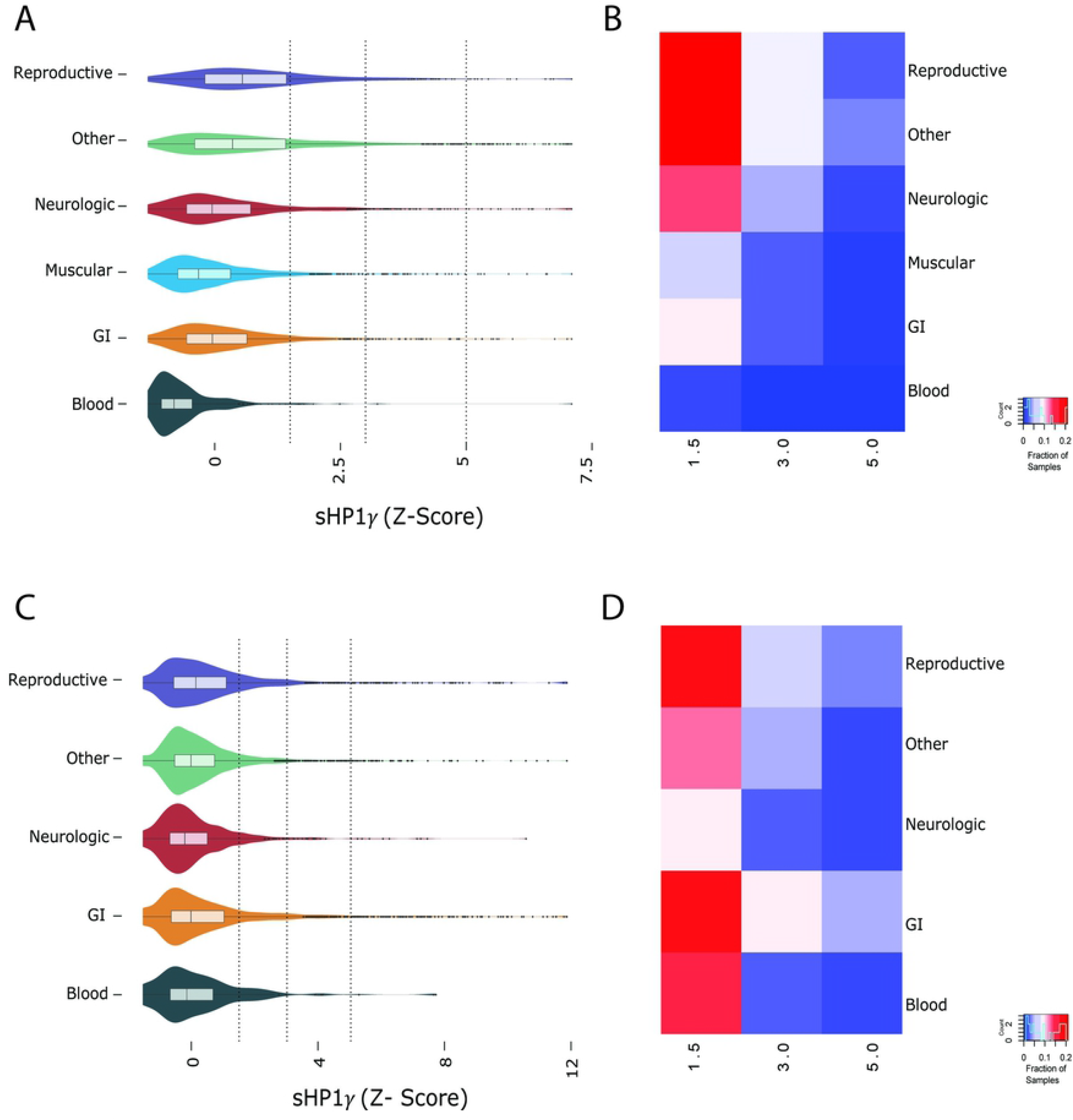
Expression level of sHP1γ is variable across normal human and cancer tissues. Gene expression data gathered from GTEx shows the (A) distribution of gene expression levels across six groups of human tissues. Some tissues have markedly higher expression of sHP1γ than others. Vertical dotted lines indicate the global 85th, 95th and 97.5th percentiles. (B) The fraction of samples from each tissue group with an expression level at least the 85th (5 Transcripts Per Million - TPM), 95th and 97.5th percentiles are shown as a heatmap. Gene expression data extracted from TCGA shows the (C) distribution of gene expression levels across five groups of human tumors. Vertical dotted lines indicate the global 85th, 95th and 97.5th percentiles. (D) The fraction of samples from each tumor group with an expression level of at least the 85th (5 TPM), 95th and 97.5th percentiles are shown as a heatmap.

### sHP1γ is translated in human cells where it localizes to the nucleus

Numerous alternatively spliced transcripts are often found in RNA-Seq studies; however, not all of these transcripts are reliably detected as alternative isoforms at the protein level[20]. To confirm that the transcript gives rise to a stable sHP1γ protein, we generated a polyclonal antibody against a synthetic peptide (RKEMLLTNQEDLPEVLILKE) for use in both, western blot (Fig 4A) and immunofluorescence (Fig 4B). The peptide was submitted to Basic Local Alignment Search Tool (BLAST, NCBI) and found to be present exclusively in the C-terminal region of sHP1γ. For the western blot analysis, we transiently transfected CHO cells with an epitope-tagged (His) form of sHP1γ or full length HP1γ, as a control, to induce expression of these isoforms. Fig 4A shows that the sHP1γ band is specifically detected in cells that overexpress the small isoform, and the antibody does not cross react with the full-length gene product. The overexpression of both isoforms was confirmed using a His tag antibody (Fig 4A). Therefore, this data demonstrates that the sHP1γ mRNA is expressed and translated into a stable protein *in vitro*.

**Fig 4.**
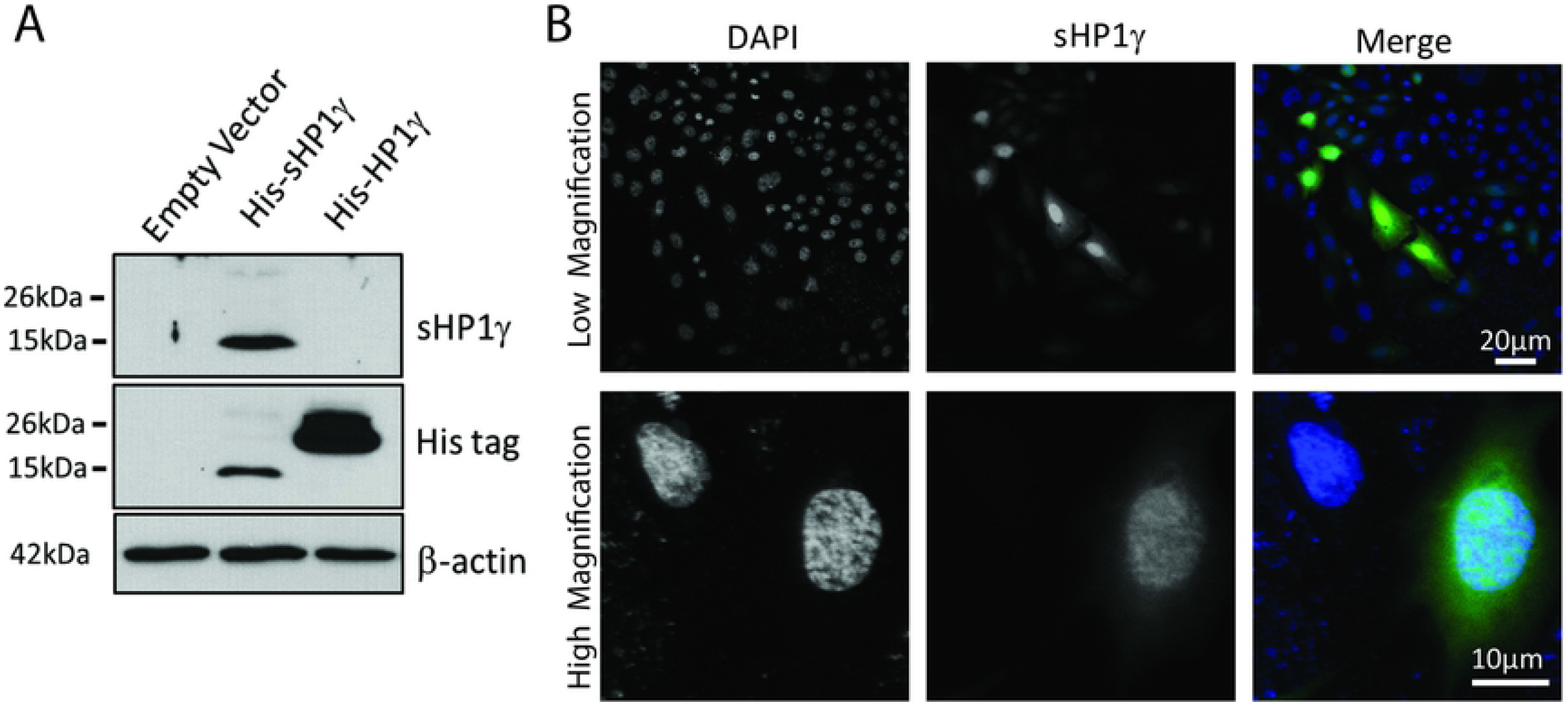
Detection of sHP1γ protein. (A) Lysates from CHO cells transfected with empty vector, His-sHP1γ and His-HP1γ were used for Western blot analyses with a newly generated peptide-specific antibody against sHP1γ. The overexpression of both HP1γ isoforms was confirmed with His antibody and β-actin was used as reference control. Molecular weight markers on the left illustrate the size difference of the two HP1γ isoforms. (B) Immunofluorescence analysis was performed for sHP1γ using our sHP1γ specific antibody (green) in HeLa cells. Independent fields of stained cells are shown at low and high magnification, upper and lower panel respectively. DAPI staining (blue) was carried out under optimum conditions to reveal nuclear structures.

Using an adenovirus construct and the same sHP1γ antibody, we transduced, fixed and conducted immunofluorescence confocal microscopy with HeLa cells to gain insight into the expression and localization of sHP1γ. In line with previous reports on the CSD-containing conventional HP1γ[21], we detected a strong signal for sHP1γ in the cell nucleus (Fig 4B), where other members of this family of chromatin proteins work. We also detected weak presence of sHP1γ in the cytoplasm, which may reflect the ability of these proteins to bind to α-importin, as demonstrated by our group, to transport HP1 proteins from the cytoplasm to the nucleus[8]. This experimental evidence is congruent with SLiM analyses, as performed by the ELM software[22], which identifies a monopartite NLS present in the non-conserved region of sHP1γ (a.a. 57-64). Interestingly, in contrast, the long canonical isoform bears a bipartite form of this motif that conforms to the KRKS-(X_9_)-KSKKKR consensus sequence. In spite of many years of investigations in the field of protein localization, no significant qualitative or quantitative differences have been reported between these two types of domains. Thus, functionally, they are both considered to be highly effective nuclear targeting sequences, which is congruent with our results that both isoforms primarily localize to the cell nucleus. In summary, the combined data from western blot and immunofluorescence analyses complement our findings with RNA-Seq and isoform-specific qPCR to reveal for the first time that the *CBX3/HP1γ* gene, which is regulated by alternative splicing, is translated to a shorter HP1γ protein in human cells where it primarily localizes to the cell nucleus. Thus, we subsequently studied the molecular properties of this novel HP1γ isoform, using sequence-based bioinformatics approaches, as well as homology-based molecular modeling and dynamic simulations.

### Modeling sHP1γ as a CSD-Less Protein with the Ability to Bind to Methylated Histone Peptides

To begin characterizing structural and molecular dynamic properties of this new sHP1γ protein, we utilized structural bioinformatics, molecular modelling, and molecular dynamic simulations. Order-to-disorder predictions indicated that both the N (1-Met to 30 -Phe) and C-terminal (97-Leu to 101-Glu) regions of sHP1γ are intrinsically disordered regions (IDR), while the residues located between them (31-Val to 96-Val) exhibit the opposite characteristics (Fig 5A). Interestingly, the final 45 amino acids at the C-terminus have no sequence similarities with other members of the CBX family, to which this protein belongs. However, fold recognition analyses using the JPRED4 algorithm[23] revealed that the sequence scores well for the CD protein fold and for protein folds with a similar organization, consisting of N-terminal β-sheets followed by an α-helix (Fig 5B). Critical assessment of structure predictions (CASP)-high scoring approaches including I-TASSER[24] and X-Raptor[25] used for sequence-to-structure predictions agree with this data (S3 Fig). Analysis sHP1γ using BLAST revealed a clear lack of the C-terminal CSD. Based on these analyses, the predicted organization of this protein is IDR- (β-sheets-α-helix)-IDR.

**Fig 5.**
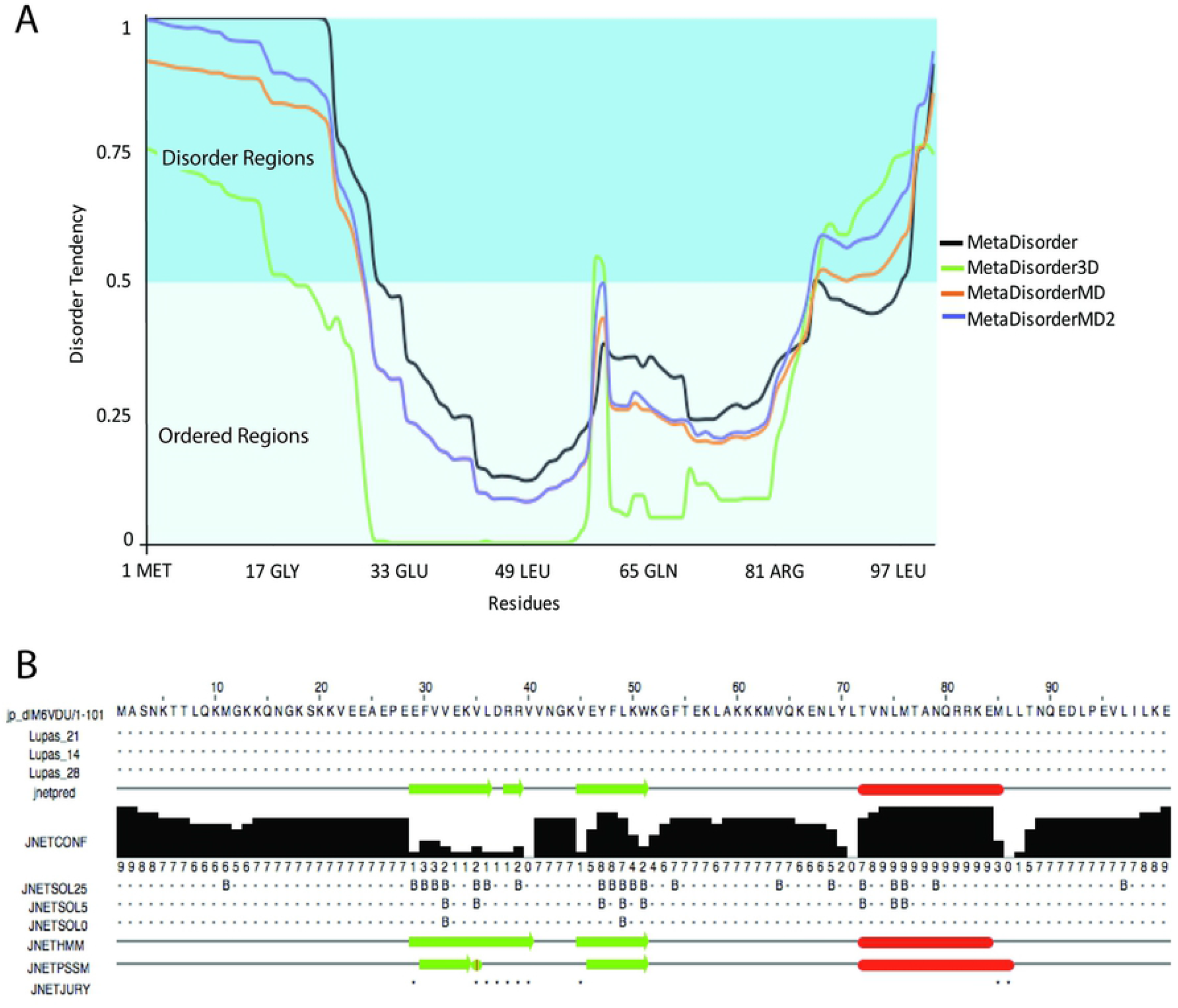
sHP1γ folding characteristics. (A) Protein disorder plot for sHP1γ generated by MetaDisorder software. All residues whose disorder probability is over 0.5 are considered as disordered. (B) The secondary structure of sHPγ generated by JPRED4. β-sheets are marked as green arrows and α-helices as red bars.

To shed light on the molecular properties of sHP1γ, we also generated a 3D model by satisfying spatial restraints deployed by MODELLER[26] and utilizing experimentally solved structures as templates (PDB IDs: 2L11, 3KUP, and 3DM1)[27, 28]. The structural model obtained for sHP1γ allows us to infer and compare characteristics of this isoform to the conventional isoform at atomic resolution and is shown in Fig 6A (sHP1γ upper, HP1γ lower). The quality of this model was high, with more than 96% of its residues being present in the allowed region of the Ramachandran plot (S4 Fig). Notably, this sHP1γ model reflects an N-terminal IDR followed by a globular domain containing the typical arrangement of an N-terminal three-stranded β-sheet that packs against a C-terminal α-helix, adjacent to a C-terminal IDR (Fig 6A). Within the conserved CD, the second and third β-strands can be connected by loops of variable lengths. However, CDs with longer or shorter linkers connecting the β-sheets to the helix are not found throughout evolution, as they would disrupt the core of structure[29]. This is consistent with the results of structural comparisons, which indicate that the CD of this human protein is conserved among isoforms and those present in evolutionarily distant organisms such as yeast, flies, and plants (Table 1). Along with these measurements of structural similarities, we show an example of a structural overlay between sHP1γ and MMP8 (Fig 6B), another CD-containing, CSD-less protein that binds to H3K9Me3, but is encoded by a non-HP1-encoding human locus[30]. Docking of the H3K9Me3 peptide to form a complex with sHP1γ was feasible through conservation of its CD with the CSD-containing conventional long HP1γ isoform (Figs 6C, 6D and 6E). Direct bonds formed between the aromatic cage of the sHP1γ CD and the H3K9Me3 mark are represented in Table 2 and S1 Table. The space between the turns and the helix defines a cavity or channel to horizontally accommodate the histone tail peptide (Fig 6C). A surface rendition of the sHP1γ-H3K9Me3 peptide complex, displayed in Fig 6D, better shows how the histone tail peptide becomes buried into this cavity. Figs 6E and 7A depict the contact between the H3K9Me3 mark and the aromatic cage formed by F30, W51, and F54. Molecular dynamics (MD) simulations (5 ns) revealed a time-dependent interaction between these aromatic residues and H3K9Me3 (Fig 7B). In addition, we noted during dynamic simulations that other residues, such as E26 and E28, are also critical for maintaining these intermolecular interactions (Fig 7B). Root mean square fluctuations (RMSF) values obtained during MD simulations, as a measure of regional displacements, demonstrated that the while the IDR and helix region of these protein are quite dynamic, binding to the histone tail restricts the fluctuations of the CD (Fig 7C). To guide protein purification experiments that consider hydrodynamic radius, such as in gel filtration or ultracentrifugation, we modeled the surface properties of sHP1γ as a globular protein in solution (Fig 7D). We measured the molecular properties of this protein when given a water-accessible surface, according to the method of Neil R. Voss and Mark Gerstein[31]. These properties, which are listed in Table 3, when compared to the conventional HP1γ as modeled by Velez, *et al*.[8] indicate that sHP1γ is significantly smaller (sHP1γ volume 16443Å^3^ as compared to HP1γ, 32517Å^3^) and more spherical (sHP1γ 0.53Ψ as compared to HP1γ 0.39Ψ). Therefore, although shorter in length and different in sequence, sHP1γ appears to conserve some of the most salient structural features of the CBX family of proteins. These findings should be taken into consideration when studying the repertoire of HP1 proteins expressed in humans.

**Fig 6.**
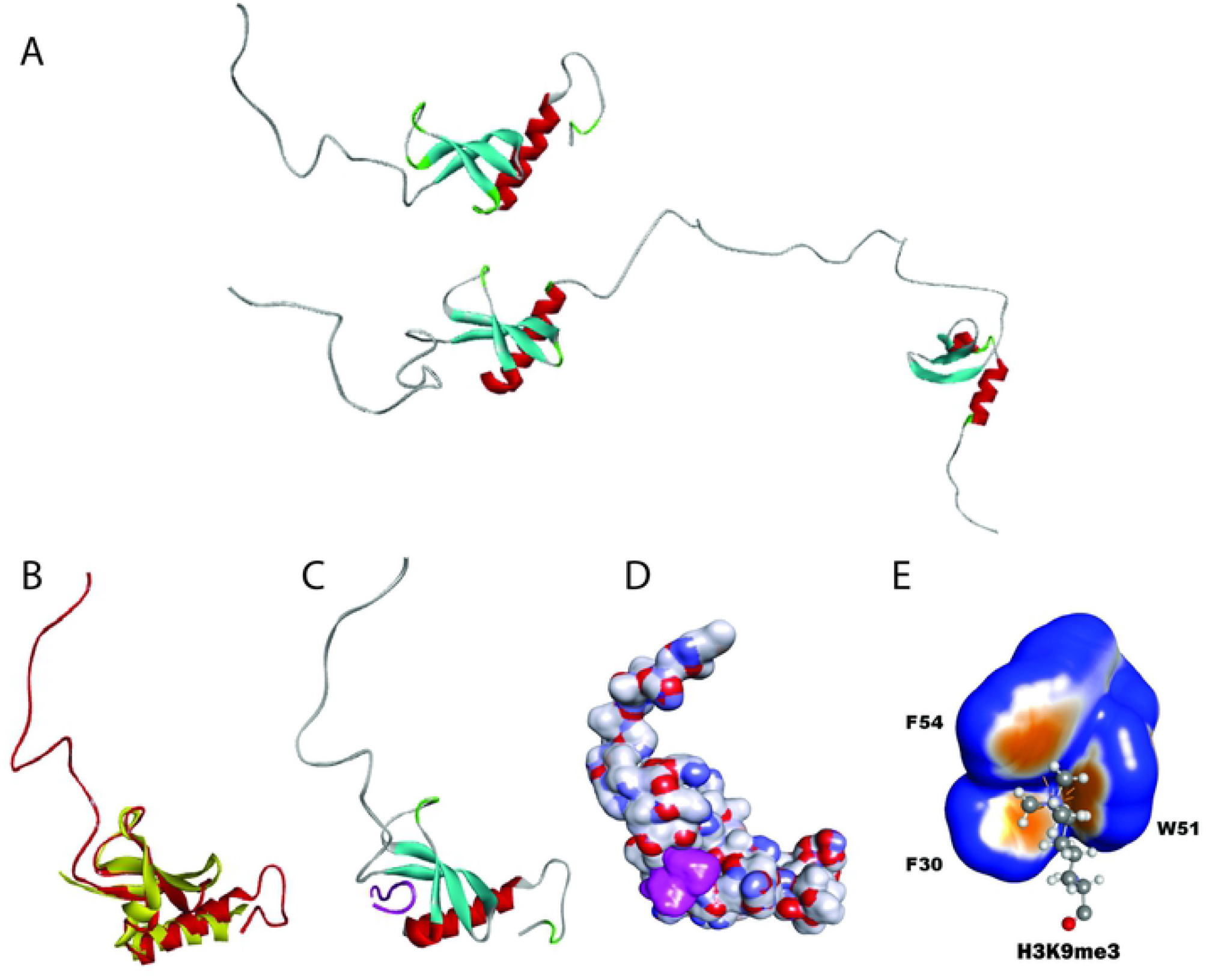
sHP1γ Comparative and Structural Molecular Modeling. (A) Molecular modeling of the novel sHP1γ (upper) and conventional full-length HP1γ (lower) isoforms in ribbon representation. The molecule can be divided into an N-terminal intrinsically disordered region (IDR; grey color), a chromodomain (β-sheets, light blue and α-helix, red), and a C-terminal IDR. (B) Structural model of sHP1γ (red) overlayed with MMP8 (yellow). (C) Structural model of sHP1γ complexed with a H3K9Me3 histone mark peptide (magenta). Tertiary structure showing how the chromoshadow-less sHP1γ accommodates a H3K9Me3 histone mark-containing peptide in a binding cavity provided by the chromodomain. The molecule can be divided into a N-terminal IDR (grey color), a Chromodomain (β-sheets colored light blue and α-helix, red). (D) Solvent accessible surface representation of sHP1γ (atom charge representation) reading the H3K9Me3 histone mark (magenta). Tertiary structure shows how the chromoshadow-less sHPγ accommodates a H3K9Me3 histone mark-containing peptide in a binding cavity provided by the chromodomain. (E) Close view of the sHP1γ aromatic cage establishing contact with K3K9M3. The light brown areas correspond to more aromatic character.

**Fig 7.**
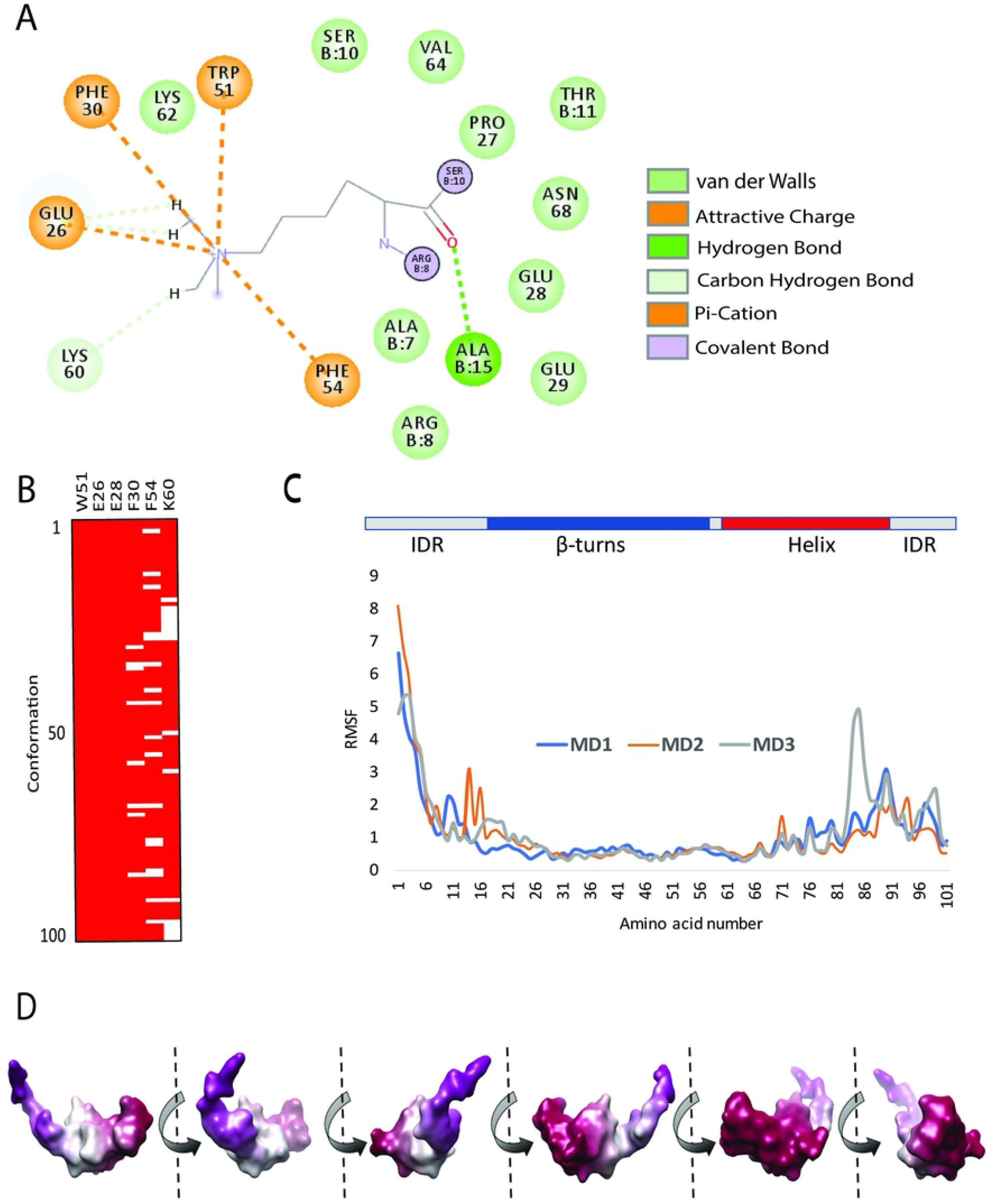
Time Dependent Interaction of sHP1γ with the H3K9Me3 Histone Mark peptide. (A) 2D diagram of the bonding pattern of H3K9Me3 to the sHP1γ aromatic cage. The diagram also shows additional stabilizing bonds (A15, E26 and K60). (B) Heat map showing that the specified amino acids maintain time-dependent contacts with the H3K9Me3 residue. Amino acids W51, E26, and E28 make contact with the H3K9Me3 residue in 100% of the conformation sampled during 5 nanosecond MD simulations. (C) The sHP1γ diagram shown at the top represents the secondary structural features of this protein. The low RMSF values corresponding to the β-turn containing regions (aromatic cage), which bind to the histone mark. (D) Surface-derived molecular properties of sHP1γ. Images correspond to all faces of which were used to determine the molecular properties listed on Table 3.

**Table 1:** Similarities of sHP1γ to Chromodomain containing proteins across kingdoms. Root mean square deviation (RMSD) values provide insight into the similarities of sHP1γ to organisms ranging from human to *saccharomyces pombe* (1e0b-B) and *drosophilae melanogaster* (1kne-A, and 5xyw-B) to even plants (4iut-A). The protein data bank (PDB) identifiers are listed for each ortholog.

| PROTEIN | PDB | RMSD |
| --- | --- | --- |
| CBX3 | 2l11-A | 0.7 |
| CBX5 | 3fdt-A | 1 |
| CBX6 | 3i90-A | 1.1 |
| CBX7 | 4mn3-A | 1.2 |
| *d.m* HP1 | 1kne-A | 1.2 |
| RHINO | 5xyw-B | 1.2 |
| *d.m* PLC | 1pdq-A | 1.3 |
| MMP8 | 3r93-C | 1.4 |
| SUV39H1 | 3mts-C | 1.4 |
| CBX8 | 3i91-A | 1.4 |
| CBX2 | 5epk-A | 1.5 |
| a.t. SAWADEE | 4iut-A | 1.5 |
| CBX1 | 3g7l-A | 1.6 |
| *s.p.* HP1 | 1e0b-B | 1.7 |
| CDYL-2 | 5jjz-A | 1.7 |

**Table 2:** Direct bonds formed between the aromatic cage of sHP1γ CD and the H3K9me3 mark.

| Bond Type | From Residues | To Residues |
| --- | --- | --- |
| Hydrogen Bond | C:M3L9:HN | A:GLU39:O |
| Hydrogen Bond | C:M3L9:HM31 | A:PHE42 |
| Electrostatic | C:M3L9:NZ | A:PHE42 |
| Electrostatic | C:M3L9:NZ | A:TRP65 |
| Electrostatic | C:M3L9:NZ | A:TRP65 |

**Table 3:** Surface-Derived Molecular Properties of sHP1γ

| Property | Unit |
| --- | --- |
| Voxel Size | 0.5 Å |
| Volume | 16443 Å<sup>3</sup> |
| Surface Area | 5942 Å<sup>2</sup> |
| Sphericity | 0.53 $\Psi$ |
| Effective Radius | 8.30 Å |
| Center of Mass | (-13.0, -4.7, 0.9) Å |

## Discussion

HP1 was one of the first epigenomic regulators identified through its function in heterochromatin formation. Thus, the current study contributes significantly to advance this field, by reporting the identification and characterization of a small HP1γ isoform. The novel isoform was discovered as a result of our long-term efforts to better understand the role of this protein family in pancreatic cancer[32, 33]. Reanalysis of RNA-Seq samples from pancreatic cancer cells[12] led us to define that the *CBX3* locus gives rise to mRNA that encodes a novel spliced form of the *CBX3* locus, which we termed small HP1γ, sHP1γ. We present both bioinformatics-based and experimentally-derived evidence demonstrating that the mRNA for this novel isoform is widely expressed in both normal and many different cancer types of human tissues. A peptide-specific antibody against sHP1γ confirmed that this protein is indeed translated. Furthermore, immunofluorescence-based microscopy corroborated bioinformatics-based predictions for this protein with localization to the cell nucleus. Thus, combined, these experiments report, for the first time, the existence of a *bona fide* CSD-less HP1γ isoform, which is not specific to PDAC, but rather widely expressed in a variety of human tissues.

To gain insight into the molecular properties of this novel isoform at an atomic level, we modeled the structure and dynamics of this protein, using similar methodologies to those we have previously applied to study properties of the full-length conventional HP1γ isoform[8]. Sequence analyses demonstrated that sHP1γ shares complete conservation of the CD region, which contains the residues that form the aromatic cage used by HP1 and Polycomb (Pc) CBX family members for binding methylated histone 3 at K9 and K27, respectively. Notably, the rest of the protein does not contain homology to the larger isoforms. A meta, order-to-disorder analysis of the CD predicted that sHP1γ will likely fold in a manner that preserves the methylated histone binding function of this domain. Structural predictions that use a combination of threading (as in Fig 5B) and homology-based (as in Fig 6) methods revealed the potential of sHP1γ to adopt a structure that is highly similar to most members of the HP1/Pc CBX family of proteins[28]. Pure homology modeling by satisfaction of special restraints using MODELLER[26] was congruent with the other methods. To gain insight into the molecular properties of this protein, we built a model of sHP1γ both, in its free form and when bound to the H3K9Me3 peptide. Molecular dynamic simulations suggested that the properties of this protein are consistent with those previously described for other family members (HP1α, HP1β, HP1γ)[8]. When compared with the conventional HP1γ isoform, however, this protein lacks a C-terminal CSD domain. On the other hand, the structure of this protein highly resembles another H3K9Me3 binding protein, MPP8, which also lacks a CSD and is encoded by a gene distinct from CBX3[30]. Thus, our study provides evidence that an expanded number of proteins can bind to H3K9Me3. These proteins can be divided into two groups: those proteins that contain a C-terminal CSD and another set of proteins which lack this domain. From this observation, we can draw inferences that are important to consider as it relates to sHP1γ. For instance, the lack of the CSD used for heterodimerization among the conventional HP1α, HP1β, and HP1γ, as well as for recruiting many other partners that regulate many cellular functions, is highly suggestive that sHP1γ will have some divergent functions. If the protein behaves more like other CSD-less CBX members[29], one could expect that it may use the CD for the dual function of binding histones and forming complexes with other proteins. For instance, the longer HP1γ isoform recruits its histone methyl transferase partner, SUVH39H1, through the PXVXL motif that is formed by its dimerized CSD[4], a function which sHP1γ would not be able to perform, at least through a similar binding mechanism. On the other hand, G9a/EHMT2 is another histone methyl transferase that is recruited by HP1γ through its CD[34], a phenomenon in which sHP1γ may also participate. Thus, we are optimistic that future studies from our laboratory focused on directly addressing this question will further contribute to define the complexity of this system.

In conclusion, we have used a combination of experimental, bioinformatics, modeling, and molecular dynamics methods in the current study to describe the existence of a novel CSD-less sHP1γ. This discovery extends the repertoire of proteins that may bind H3K9Me3 under similar or distinct functional contexts. Because of the known role of these proteins in both physiological and pathological processes, this data bears both biological and biomedical relevance. Furthermore, the insight provided here offers speculation on the regulation of pathways utilizing HP1 proteins and broadens the functional repertoire of these important epigenomic regulators for future investigations.

## Materials and Methods

### Cell Lines and Reagents

Human pancreatic cells, CHO and HeLa cells were obtained from American Type Culture Collection (ATCC) and cultured in appropriate media at 37 °C with 5% CO_2_ according to recommendations. Bruns *et al.[35]* originally isolated the L3.6 cells which were maintained in minimum essential media (Invitrogen) supplemented with 10% fetal bovine serum, 2mM L-glutamine (Gibco), 1x MEM nonessential amino acids (Gibco), 2x MEM vitamins (Gibco), 1mM sodium pyruvate (Gibco) and 0.1% antibioitic/antimycotic (Invitrogen). Standard molecular biology techniques were used to clone sHP1γ and HP1γ, as previously described[36] into pcDNA3.1-His and Ad5CMV vectors. For transient expression in CHO cells, 1×10^6 cells were plated in 60mm dishes and allowed to attach overnight. Cells were allowed to recover for 48 hours after Lipofectamine2000 (Invitrogen) transfection with 8μg of the His-HP1γ, His-sHP1γ or His-Empty Vector control plasmid. Recombinant adenovirus for sHP1γ and HP1γ[36] was generated through the Gene Transfer Vector Core at the University of Iowa, IA, USA. The generation and purification of an antibody against sHP1γ was completed using a similar method as previously described by our laboratory for the canonical isoform[16] using the immunogen peptide with the RKEMLLTNQEDLPEVLILKE primary sequence.

### Identification and Tissue Distribution of the sHP1γ Encoding mRNA

Evaluation of the normal human tissue panel was performed using FirstChoice® Human total RNA Survey Panel following manufacturer’s recommendations. Total RNA for stomach and pancreas tissues were purchased separately from Agilent/Stratagene (Catalog# 540023 pancreas and 5400037 stomach). For human pancreatic cancer cell lines, total RNA was isolated with the Qiagen RNeasy kit according to the manufacturer’s protocol. RNA was reverse-transcribed using the SuperScript III System (Invitrogen), and SYBR Green-based real-time PCR was performed according to the manufacturer’s instructions (Bio-Rad). Primers for isoform-specific qPCR were designed with the following sequences: sHP1γ F: 5-GTGGAAGGGATTTACAGAAAAGC-3, sHP1γ R: 5-TTTGCCAGAGGTCTTGATCC-3, HP1γ F: 5-AAGGGATTTACAGATGCTGAC-3 and HP1γ R: GACAAACCAAGAGGATTTGC.

### Western blot analysis

For Western blots, transfected CHO cells were lysed in 4× Laemmli buffer (250 mm Tris (pH 6.8), 20% glycerol, 8% SDS, 0.0025% bromophenol blue, 1 mm *β*-mercaptoethanol), collected by scraping with a rubber policeman, sonicated for 10s, and boiled at 95 °C for 5 min prior to loading. Lysates were subjected to 14% SDS polyacrylamide gel electrophoresis, transferred onto polyvinylidene difluoride membranes and probed with the anti-sHP1γ antibody diluted in 5% milk in tris buffered saline with tween (TBST). Anti-rabbit secondary antibodies (Millipore) were incubated on the membranes for 1 h at room temperature, after three successive washes with TBST, bands were detected with enhanced chemiluminescence (ECL, Pierce).

### Immunofluorescence

HeLa cells were plated on poly-L-lysine-coated circular coverslips (0.1 mg/mL poly-L-lysine) and allowed to adhere overnight. Transduction occurred at a multiplicity of infection of 400:1. After 48 h, cells were fixed with 4% formaldehyde, permeabilized with 0.2% Triton-X 100, blocked for 30 min, and finally stained with the anti-sHP1γ antibody (1:250) and Alexa Fluor 488 anti-rabbit secondary antibody (1:500, Invitrogen). Coverslips were mounted in VectaShield mounting media with DAPI for immunofluorescence. Images were acquired using 40× and 100× objective lenses on a Zeiss LSM 780 confocal microscope.

### Molecular Modeling and Molecular Dynamic Simulations

The modeling for the sHP1γ alone and in complex with the H3K9Me3 peptide was performed using a combination of approaches previously described[8]. Briefly, homology-based modeling was performed using MODELLER[26] with previously solved structures as templates (PDB IDs: 2L11, 3KUP, and 3DM1)[27, 28]. The model was refined by energy minimization using two cycles of 200 steps of steepest descent and two cycles of Random Newton before proper evaluation of stereochemical properties by the Ramachandran method. Molecular dynamic simulations were performed for 5 nanoseconds, each using different random seeds, an isothermal-isobaric (NPT) ensemble, and the distance-dependent dielectric method for simulating implicit solvent conditions. Bonds were modelled by a distance dependence approach. Since it is known that the chromodomain-H3K9Me3 complex forms by an induced fitting mechanism, a soft harmonic restraint, using the best fit intermolecular interaction method, was applied to the structure. Visualization and illustration were done using Discovery Studio (Biova).

## Abbreviations

HP1: (Heterochromatin protein 1),
CBX3: (Chromobox containing protein 3),
IDR: (intrinsically disorder region),
CD: (chromodomain),
CSD: (chromoshadow domain).

## Acknowledgements

This work was supported by National Institutes of Health Grants R01 CA178627 (to G.L.) and R01 DK52913 (to R.U.), the Advancing a Healthier Wisconsin Endowment (to each G.L. and R.U.) and The Linda T. and John A. Mellowes Endowed Innovation and Discovery Fund (to R.U.).

